# A comparison of scalable approaches for the pairwise analysis of large pathogen genomic and spatial datasets: an application to studying *Mycobacterium tuberculosis* transmission

**DOI:** 10.64898/2026.05.21.726848

**Authors:** Yu Lan, Chieh-Yin Wu, Hsien-Ho Lin, Ted Cohen, Joshua L. Warren

## Abstract

Pairwise analysis of genomic and spatial data offers opportunities to identify and estimate the associations between covariates and the transmission of pathogens between individuals. However, such pairwise analyses are computationally intensive, and may not be feasible to conduct given the high dyad count in even moderately sized datasets. Here we compare two approaches to increase the efficiency of pairwise analysis for large datasets. We quantify and compare the performance of divide-and-conquer Bayesian model fitting and pairwise case-control approaches for estimating associations between individual- and pair-level covariates and shared membership in a transmission cluster. We utilize a large dataset (n=4,154) of spatially-referenced, genomically-sequenced *Mycobacterium tuberculosis* isolates collected from a single city for this analysis. We find that the case-control approach produces unbiased estimates of effect sizes with expected credible interval coverage and is more robust than the divide-and-conquer method when effect sizes are large. Thus, we recommend using the case-control approach with at least three controls per case to downscale datasets for pairwise analysis when analysis of the entire dataset is not possible. This approach mitigates the computational challenges of pairwise Bayesian modeling on datasets that require significant computational resources while maintaining desired inferential properties.

**Author Summary:** Pairwise analyses of large datasets to study pathogen transmission are computationally demanding because they typically require simultaneous analysis of each possible pair of individuals in a dataset; as datasets become larger these analyses often are not feasible to conduct even with access to high-performance computing resources. In this work, we compare a case-control approach and divide-and-conquer approaches for more efficient pairwise analysis of large datasets. Using a large dataset of *Mycobacterium tuberculosis* isolates including genetic and spatial data, we investigate the performance of each method for estimating the associations between host covariates and genetic clustering of isolates. We find that the case-control approach is generally preferred over methods which first divide the data into subsets and then combine results. While additional extensions of these analyses are needed to test the generality of these findings to other data settings, this work provides a practical way forward for the pairwise analysis of large datasets to study pathogen transmission.

## 1. Introduction

Tuberculosis (TB) is caused by *Mycobacterium tuberculosis* (Mtb), a bacterium transmitted from person to person through the respiratory route. Whole genome sequencing (WGS) can be used to identify possible instances of person-to-person Mtb transmission based on the genetic relatedness of sequenced isolates [1]. Many Mtb studies have applied a threshold for the maximum number of genetic differences (i.e., single nucleotide polymorphisms or SNPs) between isolates to identify possible transmission linkages [2], and other specialized approaches (e.g., TransPhylo [3]; Outbreaker [4]) estimate the probability of transmission between individuals based on genetic distances as well as other factors related to the accumulation of genetic differences between cases (e.g., timing of diagnoses, bacterial mutation rates). With the increased availability of high resolution genomic and spatial data, a growing number of studies have also combined spatial data with genomic relatedness to study Mtb transmission in communities [5].

Network-based regression modeling of spatially-referenced pairwise genetic similarity measures offers a novel opportunity to understand which factors are associated with Mtb transmission, while accounting for multiple sources of correlation (e.g., network dependence and spatial correlation) [6]. These models are designed to associate pairwise measures of genetic similarity (e.g., SNP distances or shared cluster membership) between two individuals with individual- and pair-specific covariates of interest. However, as the number of individuals in a study increases, the number of unique pairs increases dramatically, leading to extreme computational demands. For a dataset with n>1 individuals, there are n(n-1)/2 dyadic observations when directionality of transmission is not considered; this results in computational challenges particularly when working in the Bayesian setting.

Employment of subsampling strategies with high-performance computing (HPC) resources is one possible approach to address this computational challenge. One promising sampling strategy is the case-control approach [7, 8]. Furthermore, some studies have discussed different sampling designs, such as a two-step case-control sampling scheme for studying binary outcomes [9], optimal sampling for large sample logistic regression [10], and sampling with replacement and Poisson sampling in optimal subsampling [11].

Another computational strategy is the divide-and-conquer approach, where the large dataset is divided into subsets, analysis is conducted on each subset, and the posterior results are combined across all analyses [12]. Several methods have been developed based on this strategy to handle the analysis of large geostatistical datasets, such as spatial meta-kriging for point data [13] and scalable Bayesian modelling on areal (lattice) count data [14]. Regardless of types of data, methods have been developed to approximate posteriors samples from analyses simultaneously run on subsets of the entire dataset. The Consensus Monte Carlo method [15] approximates the complete posterior distribution by aggregating weighted averages of subposterior samples from independent Markov chain Monte Carlo (MCMC) runs. Alternatively, the semiparametric density product estimator method (DPE) estimates the subposterior density using kernel density estimation and thus combines estimated densities of subposteriors to approximate the full posterior density [16]. Despite extensive discussions in the existing literature, few studies have focused on the applications of these approaches to dyadic data.

In Section 2, we introduce and conduct a pairwise analysis on two large (n= 700) benchmark datasets with different pairwise clustering proportions from the full dataset collected in Kaohsiung, Taiwan. These two datasets were near the maximum size that we could analyze given our computational resources. We perform a pairwise analysis on these datasets to obtain “ground truth” estimates which we use to benchmark and compare estimates derived from competing methods described in Section 3. Section 3 describes the simulation study which we use to compare competing methods for obtaining estimates from subsampled datasets. Based on the results from the simulation study, we apply the top performing methods to the entire Mtb dataset from Kaohsiung (n= 4,154) and report the results in Section 4. We conclude with recommendations for pairwise analysis using Bayesian models for large dyadic datasets in Section 5.

## 2. Benchmark data and application

### 2.1 Benchmark Data

We use a population-based dataset of 4,154 individuals with culture-confirmed TB in Kaohsiung from 2019 to 2023 (Fig 1), with linked data on genomic cluster, age, and geolocation of residence for each individual. Additional details of the study from which these individuals were identified have been previously published [17] The size for our benchmark subsamples (i.e., n=700) is close to the upper limit of pairwise analysis that we can process given our available computational resources. We perform this pairwise analysis on a partition of HPC cluster with 200 GB of memory that supports jobs up to seven days in duration.

**Fig 1.**
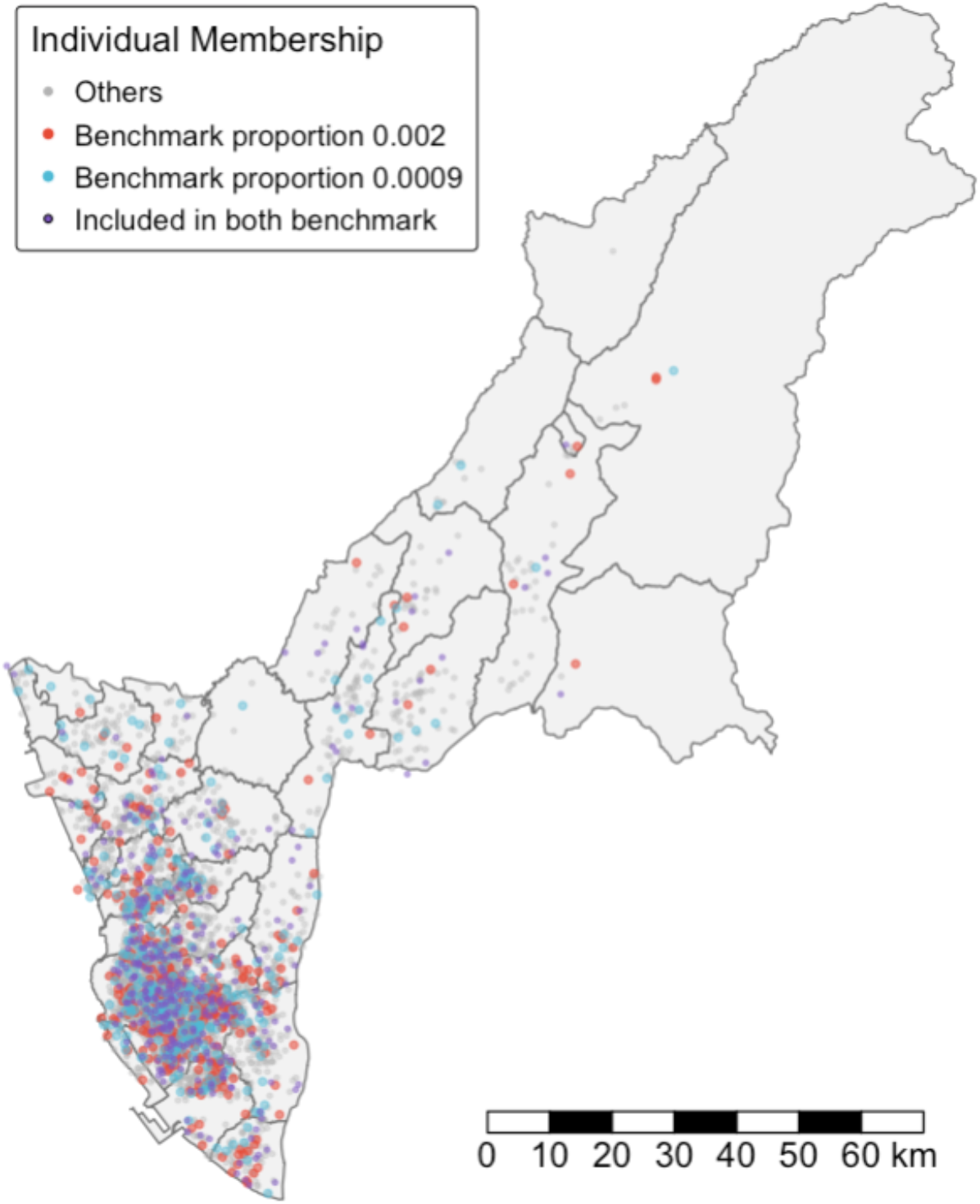
Spatial distribution of individuals with Mtb in Kaohsiung, Taiwan (2019–2023), jittered for privacy. The colored points show two sets of 700 isolates sampled from the full dataset to create the benchmark estimates with pairwise clustering proportions of 0.002 and 0.0009.

In this study, we define genomic clusters using a 12 SNP threshold. We assign pairs of individuals belonging to the same genomic cluster as having a binary shared cluster membership outcome equal to 1; all other pairs are assigned an outcome of 0. To capture varying proportions of genetic clustering in datasets, we construct two benchmark datasets characterized by different pairwise clustering proportions, defined as the fraction of pairs where the binary outcome is equal to 1 among the total number of unique pairs. Each dataset consists of 700 isolates with clustering proportions of 0.002 and 0.0009, selected from the full dataset (n=4,154). We note that the clustering proportion of the full dataset is 0.0006.

To obtain a dataset with a higher clustering proportion, we include all isolates from small and medium-sized clusters (<5 isolates, n= 606) and randomly selected an additional 94 isolates from the remaining population, yielding a clustering fraction of 0.002. To obtain a dataset with a lower clustering proportion, we include all isolates from small clusters (<3 isolates; n = 346) and randomly selected an additional 354 isolates from the remaining population, yielding a clustering fraction of 0.0009. Because clustering proportions will vary substantially across studies, we consider these two values to represent two plausible and meaningfully different clustering scenarios grounded in our data. Although both values are small in absolute terms, 0.002 is more than twice 0.0009 and, in the context of TB, represents a relatively high level of clustering.

These two sets of 700 individuals comprise 244,650 unique pairs; 487 and 213 of these unique dyads are linked within a Mtb genomic cluster (see Table 1). We consider only the unique pairs among individuals given that our goal is to determine whether they are infected with very similar isolates of Mtb (and thus potentially members of the same transmission chain), rather than to infer the direction of transmission.

**Table 1.** Genomic cluster sizes and counts of pairs for two benchmark datasets.

| Clustering proportion | Cluster size | Numbers of clusters | Unique pairs per cluster | Total unique pairs |
| --- | --- | --- | --- | --- |
| **0.002** | 2 | 175 | 1 | 175 |
|  | 3 | 52 | 3 | 156 |
|  | 4 | 26 | 6 | 156 |
|  | **Total** | **253** | **-** | **487** |
| **0.0009** | 2 | 181 | 1 | 181 |
|  | 3 | 2 | 3 | 5 |
|  | 4 | 2 | 6 | 12 |
|  | 6 | 1 | 15 | 15 |
|  | **Total** | **186** | **-** | **213** |

### 2.2 Pairwise analysis

We first use the *GenePair* R package [6] to conduct a pairwise analysis on the benchmark datasets. The primary analysis uses a hierarchical Bayesian logistic regression framework that accounts for network dependence and spatial correlation to assess the association between pairwise and individual-level risk factors and transmission. The model included three pairwise covariates: the combined age of the two individuals, the absolute age difference of the two individuals, and the spatial distance between the two individuals’ residential locations.

The logistic regression model is given as

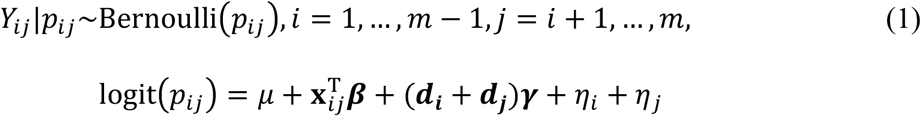

where *Y*_*ij*_ is the binary outcome describing if individuals *i* and *j* are in the same genomic cluster (1) or not (0); **x**_*ij*_ is a vector of the covariates describing the difference (i.e., the age difference and the spatial distance); (***d***_***i***_ + ***d***_***j***_) represents the covariate specific for individual *i* and individual *j* (i.e., the combined age); *m* is the number of individuals; *η*_*i*_ is the individual-specific random effect parameter modeled as a function of separable independent and spatially-correlated components, with the independent component modeled using a Gaussian distribution with mean equal to zero and unknown variance, and the spatial component modeled using a Gaussian process with exponential spatial correlation structure. Full details are given in Warren et al. [6]. The model is fitted in the Bayesian setting using MCMC sampling techniques. For each parameter, we computed posterior means and 95% highest posterior density credible intervals on the odds ratio scale.

### 2.3 Results

Analysis of the two benchmark datasets reveals that smaller age difference, a younger combined age, and shorter spatial distance between the two individuals with Mtb were independently associated with being members of the same genomic cluster (see Fig 2). We treat these parameter estimates as the ground truth for comparing the performances of the different competing computational methods.

**Fig 2.**
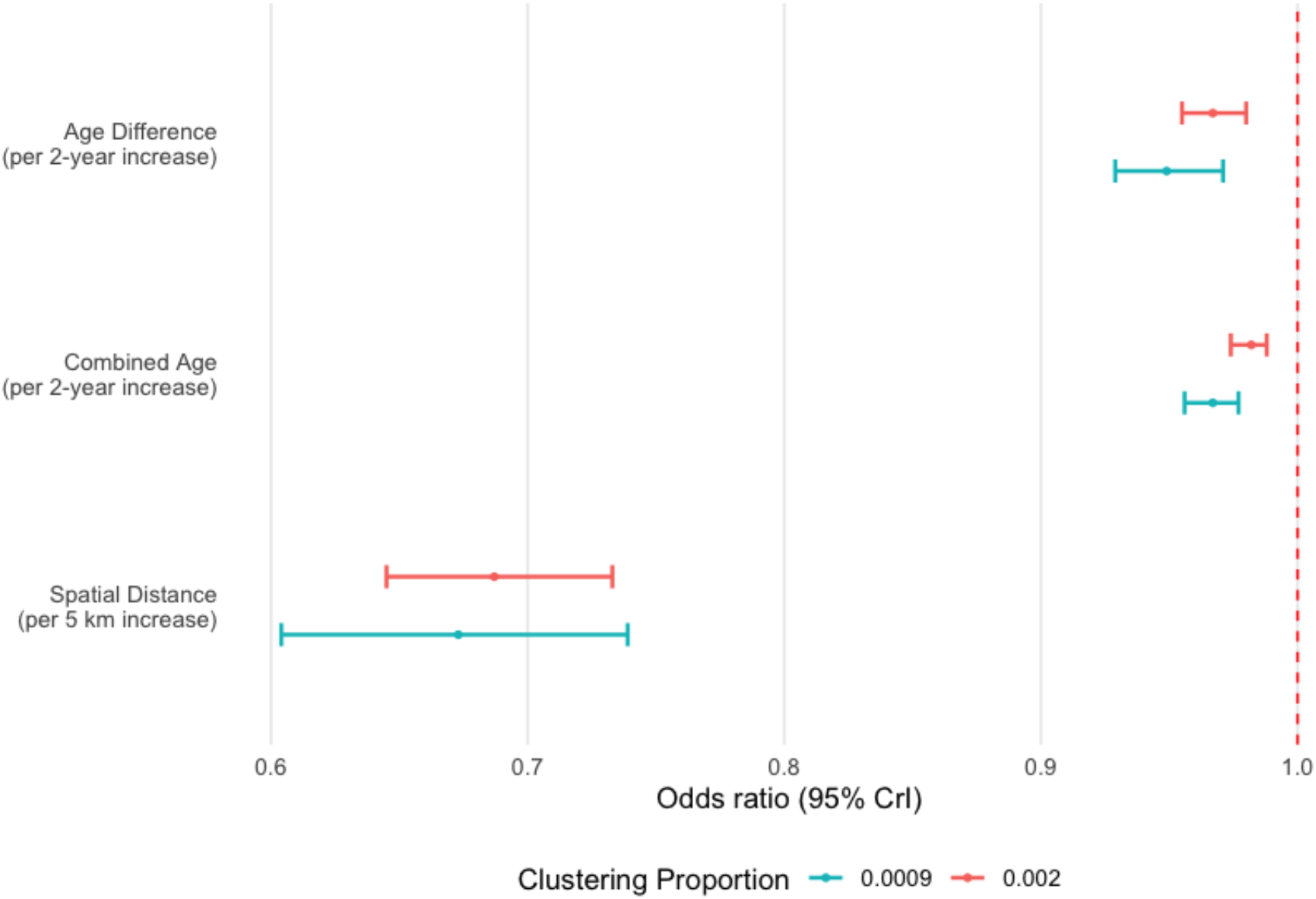
Odds ratio posterior means and 95% credible intervals for predictors in two benchmark datasets at different clustering proportion.

## 3. Comparative analysis using simulated data

### 3.1 Data generation

We simulate data from the model in (1) to evaluate the performance of the case-control and the divide-and-conquer approaches, measured by the bias of point estimates and coverage of credible intervals. The data generation process is illustrated in Fig 3. We first estimate parameters based on the GenePair results from two benchmark datasets with 700 individuals. Then, we randomly select 700 new individuals from the full dataset of 4,154 individuals for simulation purposes. Using new individuals for each simulated dataset ensures that our results are more generalizable to the full Kaohsiung dataset.

**Fig 3.**
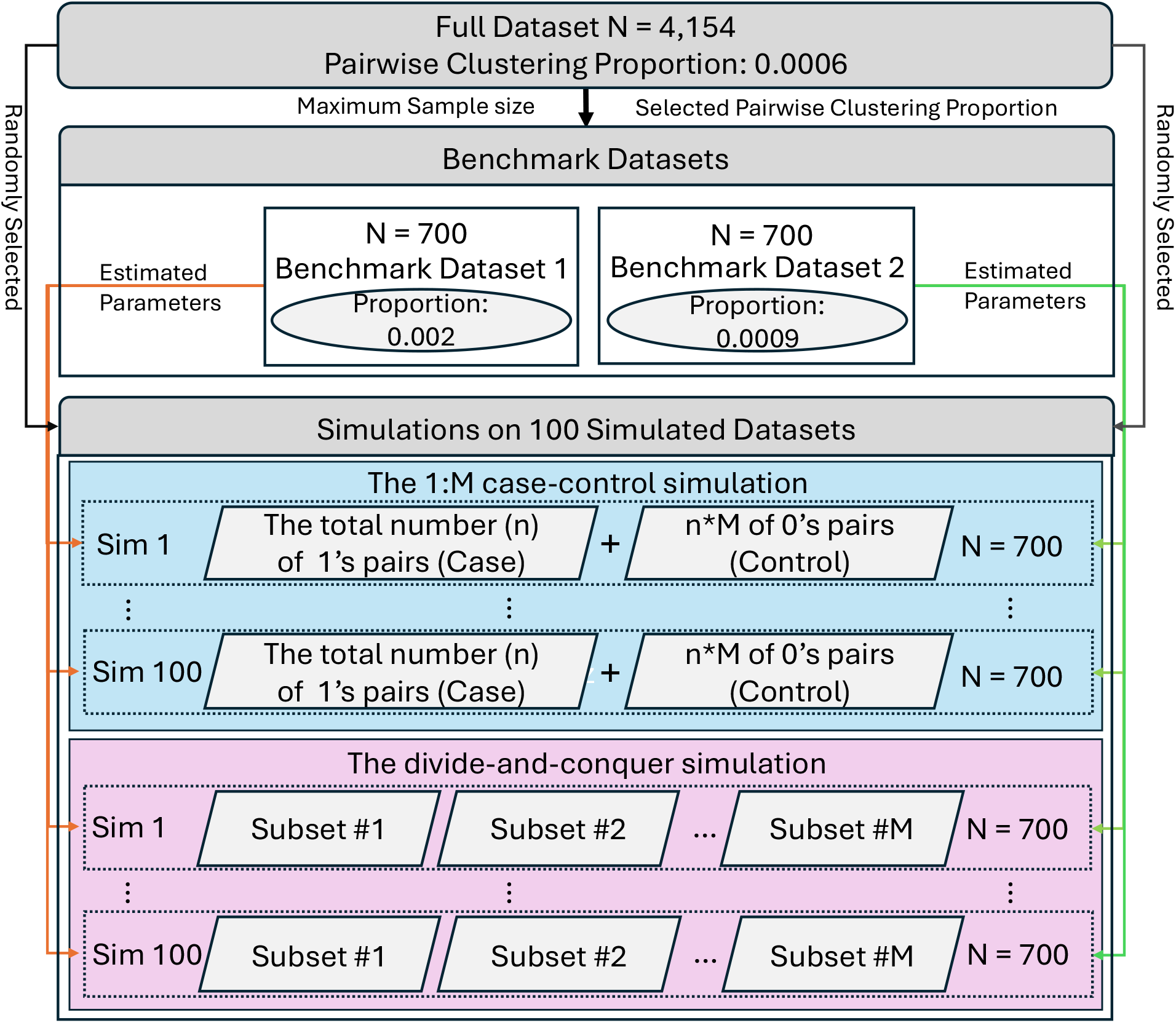
Data generation process for simulated datasets.

For each unique pair of isolates, we use the posterior mean parameter estimates from the benchmark analyses to simulate the binary outcome (*Y*_*ij*_) (i.e., whether two isolates from per the subset belonged to the same genomic cluster) using the observed covariate values for the individuals. For each simulated dataset (n=700), we generate a new realization of the individual-specific random effect parameters to avoid conditioning on a single set that ultimately define the level of correlation in the data. To do so, we use posterior mean estimates of the correlation parameters to simulate new values from both the non-spatial and spatial components for each dataset. For each clustering proportion, we simulate 100 datasets for analysis.

### 3.2 Competing Methods

To evaluate the impact of the number of controls selected per case, we compare the results when increasing the number of randomly selected controls (M_control_ = 1, 3, 5, and 10) per case. We repeat the control selection procedure for each of the four values of M_control_ on the 100 simulated datasets and run GenePair on each simulated dataset.

We investigate the divide-and-conquer strategy as follows: We divide the dataset into subsets, conduct the pairwise analysis on each subset, and combine posterior results using different statistical methods across subsets. To evaluate the impact of subset number (and therefore subset size), we conduct GenePair analysis on three values of subsets (M_subset_ = 3, 5, and 10). We repeat the subsampling procedure for each of the three values of M_subset_ on the 100 simulated datasets and ran GenePair on each simulated dataset. We use six combination methods to combine the posterior results from subsets.

We denote ***Z*** = (***Y*, x**) as the full dataset of responses and predictors. For the Bayesian pairwise analysis, the marginal posterior distribution for the regression parameters is denoted as *ρ*(***β***^*^|***Z***), where ***β***^*^ is the vector of regression parameters corresponding to the previously mentioned covariates (i.e., combined age, age difference, and spatial distance).

For the divide-and-conquer strategy, we divide the dataset into *M*_*subset*_ non-overlapping subsets ***z***_*m*_, *m* = 1, … *M*_*subset*_. We denote ***β***^*^ as the d-dimensional parameter vector of interest, and 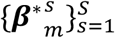 as the *s* posterior samples of the parameter vector ***β***^*^ obtained from subset *m*, where *s* = 1, … *S* and *m* = 1, … *M*_*subset*_ (after ensuring convergence). We evaluate different methods that combine the posterior samples 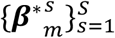 across subsets to approximate samples from the full dataset posterior distribution *ρ*(***β***^*^|***Z***). We conduct GenePair analysis on each subset and obtain MCMC samples from the posterior distribution of parameters. The six divide-and-conquer methods we include are:

- Meta-analysis (Meta) using a fixed-effect model: The final point estimate 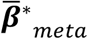 is computed as 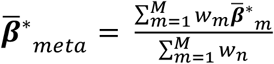, where 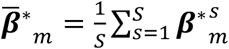, is the posterior mean estimate of the vector for subset *m*, and *w*_*m*_ is a vector of the inverse of the sample variances for each regression parameter for subset *m*.
- Sample average (sampleAvg): Approximate samples from the full data posterior distribution of ***β***^*^ by averaging the posterior samples from all subsets at each sample *s* (i.e., 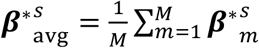).
- The Consensus Monte Carlo method: Combines independent posterior samples across subsets into pooled posterior samples. For each MCMC iteration *s*, the combined posterior samples is given by: 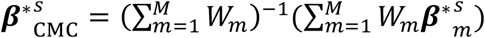, where 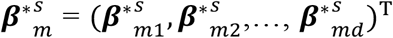 is the *s*-th posterior sample of ***β***^*^ from subset *m*, and *W*_*m*_, denotes the inverse of the covariance matrix from subset *m*. The definition of the weight matrix *W*_*m*_ depends on the assumed structure of ***β***^*^:
- MCindep: Assumes independence among model parameters ***β***^*^. Thus, *W*_*m*_ is a diagonal matrix, where each diagonal entry is the inverse of the sample variance of covariate 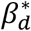, and *W*_*mj*_ is the inverse of the variance of the marginal posterior samples collected for parameter *j* after analyzing subset *m*.
- MCcov: Assumes covariance among model parameters ***β***^*^. In this case, *W*_*m*_ is defined as the inverse of the sample variance-covariance matrix estimated from subset *m*.
- DPE: Unlike other methods that directly combine posterior samples, DPE does not produce combined posterior samples directly. The full-data posterior is approximated by: 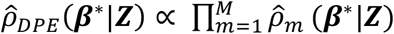, where 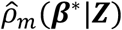 denotes the estimated subposterior density for subset *m*, obtained as the product of a parametric density and a nonparametric correction estimated via kernel smoothing. We calculate the bandwidth matrix used in kernel density estimation based on Silverman’s rule of thumb [18]. We include two bandwidth parameters in our analysis:
- DPE1 employes the original bandwidth matrix.
- DPE2 scales the original bandwidth matrix by a factor of 2 to apply greater smoothing.

We apply Meta using the *metafor* package in R [19]. We apply the other MCMC posterior combined methods (i.e., sampleAvg, MCindep, MCcov, DPE1, DPE2) using R *parallelMCMCcombine* package [20].

For each strategy, we assess bias (i.e., the difference between the estimated mean per simulation and the ground truth), the proportion of simulation replicates in which the CrI contains the ground truth, and the average 95% CrI width for each covariate individually, including the combined age, the age difference, and spatial distance.

### 3.3 Simulation results and comparison

Fig 4 shows the results for the case-control strategy using different controls per case based on all 100 simulations. Bias is shown as boxplots of the simulation-based bias estimates, together with the mean bias and its 95% confidence interval. Coverage is shown as the estimated coverage rate with its 95% confidence interval. Overall, the case-control strategy performs well with low bias and expected 95% CrI coverage for all regression parameters. For each of the two clustering proportions tested, the method with one control per case performed slightly worse than when we increased the numbers of controls. Across each clustering proportion, increasing the number of controls per case led to narrower 95% credible intervals for all three factors, indicating improved precision (see Fig S2).

**Fig 4.**
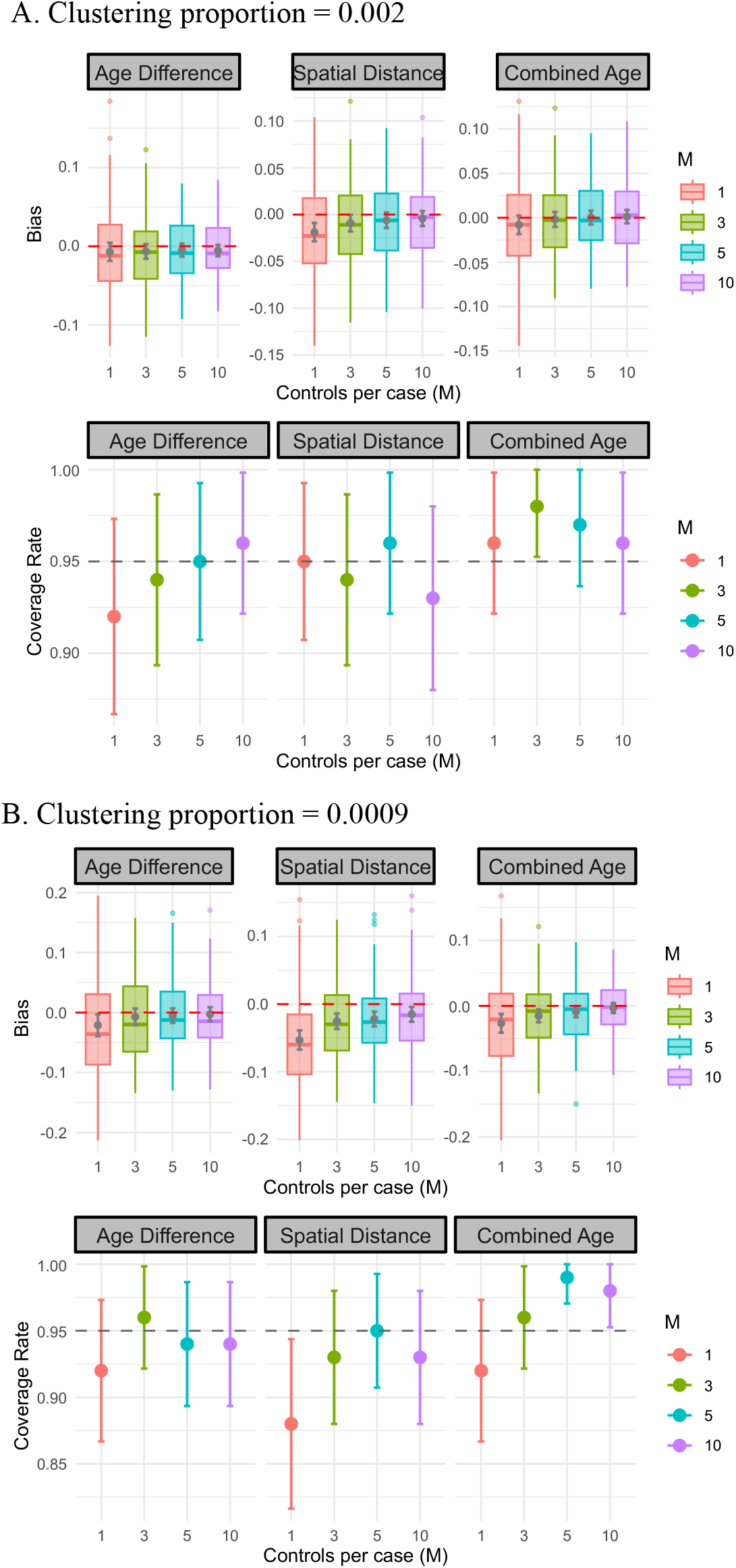
Estimated bias and empirical 95% credible interval coverage simulation study results by factors and methods using the case-control approach. Error bars show 95% confidence intervals for the estimated bias and coverage.

Fig 5 shows the results for each method using the divide-and-conquer strategy, excluding the sampleAvg method as this method has larger bias and wider CrIs than the other methods tested (see Fig S2). Overall, each method with three or five subsets performs best, and Meta was consistently among the methods that performed best with small bias and high coverage across three covariates and different types of partitions. For each clustering proportion, the divide-and-conquer approach shows wider 95% credible intervals as the number of subsets increases, indicating reduced precision as we increase the number of partitions (see Fig S3).

**Fig 5.**
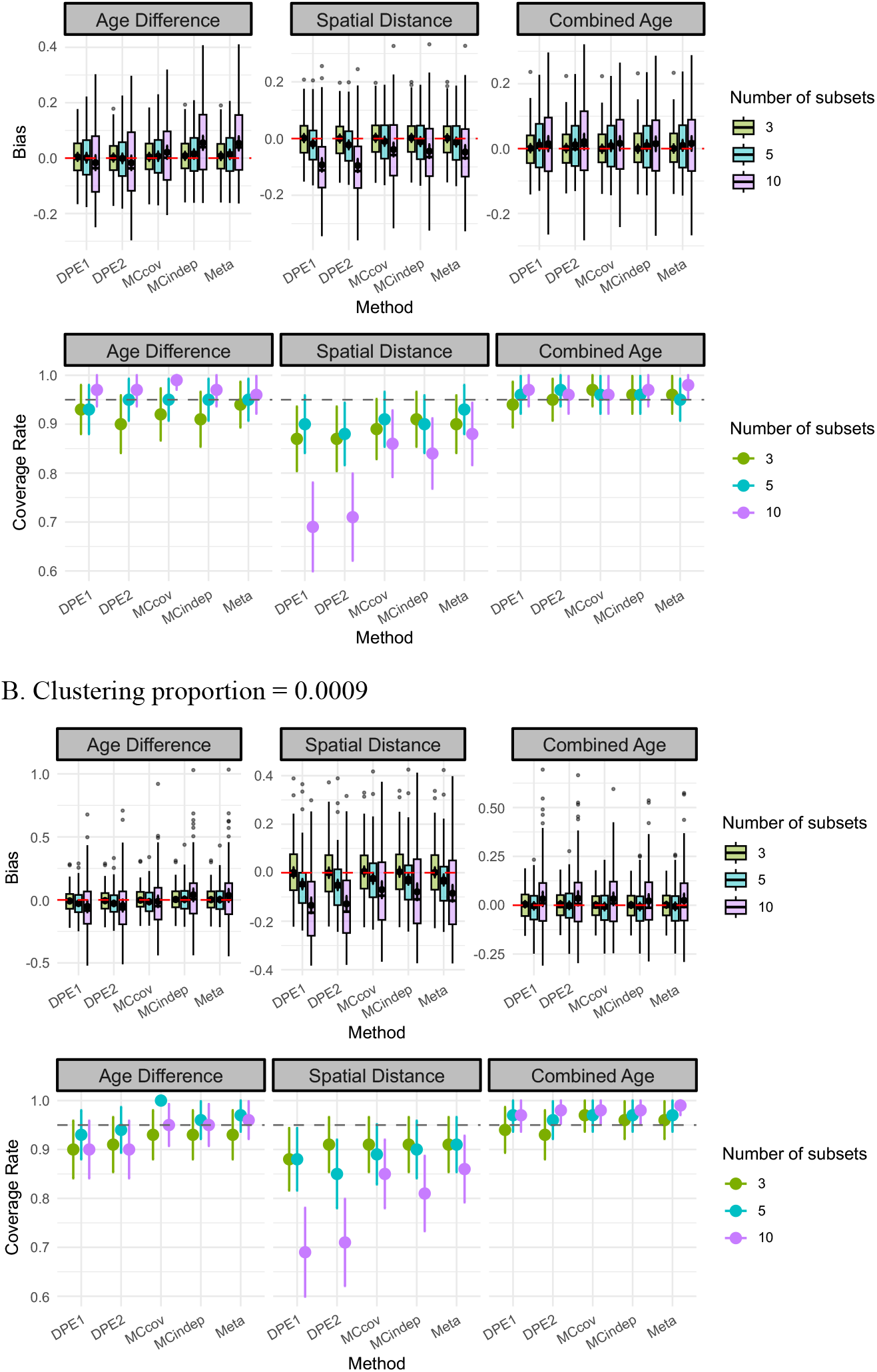
Estimated bias and empirical 95% credible interval coverage simulation study results by factors and methods using the divide-and-conquer approach. Error bars show 95% confidence intervals for the estimated bias and coverage.

Notably, in our analyses the spatial distance covariate consistently has greater bias and lower coverage compared with the non-spatial factors. When we divide the full dataset into ten subsets, the coverage rate of DPE methods decreases to approximately 70% for this variable. As shown in Fig 2, the spatial covariate is associated with the largest effect size, suggesting that this issue may not be unique to the spatial covariate, but may instead be most apparent for covariates with large effect sizes. To test this hypothesis, we swapped the effect sizes of age difference and spatial distance in the simulated datasets and observed that the poorer performance shifted to age difference (see Fig S4). This suggests that the relatively poor performance under the divide-and-conquer approach is driven by large effect sizes under sparse clustered pair information, rather than being a result of the spatial nature of this covariate.

In Fig 6, we compare the two best performing methods (i.e., lowest bias and highest coverage) from each strategy: the case-control approach with five controls per case and the divide-and-conquer approach with three subsets using the Meta method to combine subposteriors. Both methods show mean bias close to zero and similar distribution for the bias, although the divide-and-conquer method shows a wider range and a greater number of outliers. Both methods show expected coverage for the two age-related factors, while the case-control approach shows a better coverage rate for the spatial distance factor than the divide-and-conquer method. We also compare credible interval lengths of each factor estimated by the two approaches; the case–control method produces consistently narrower intervals than the divide-and-conquer method, suggesting greater precision (i.e., less posterior uncertainty) (see Fig S5).

**Fig 6.**
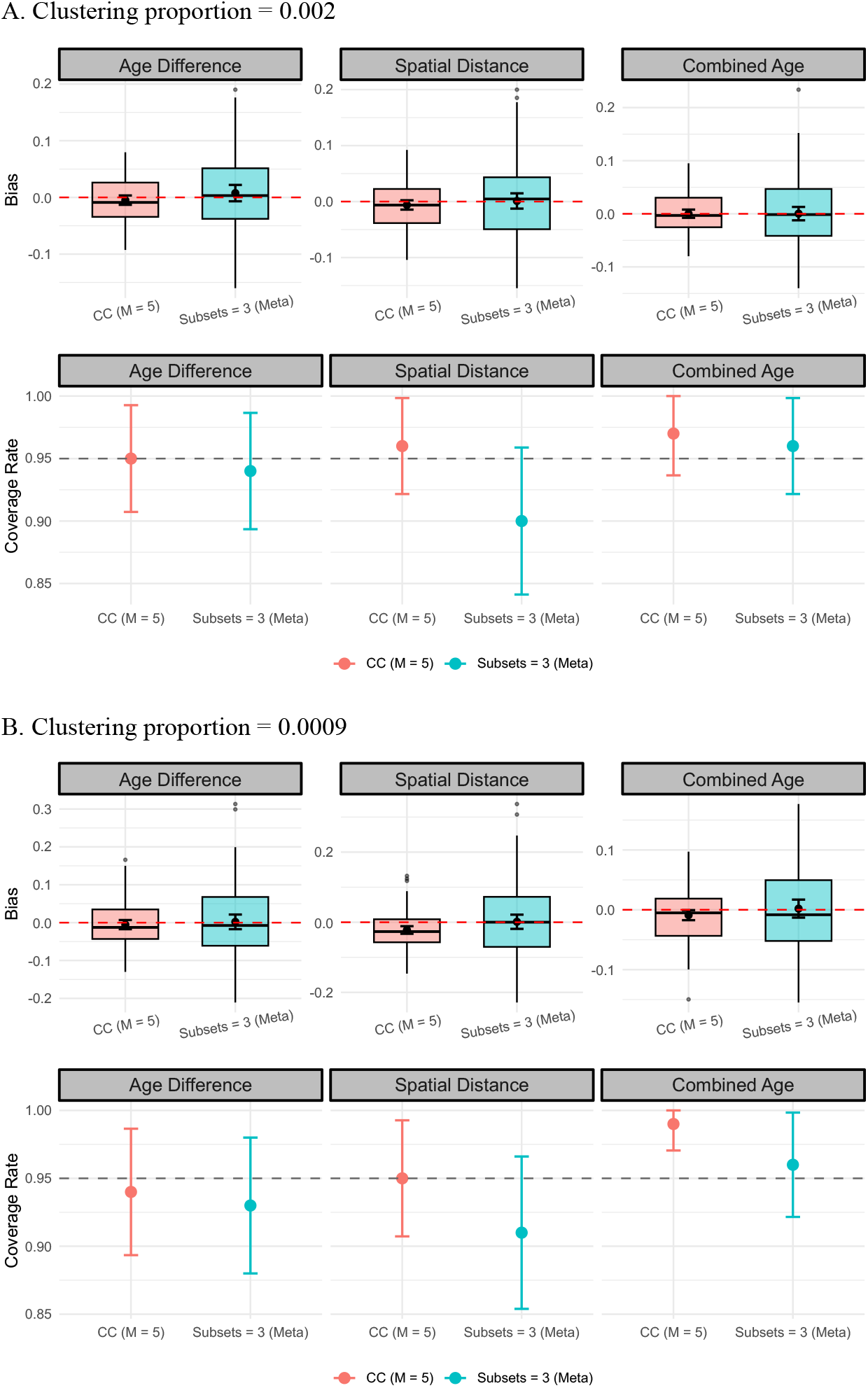
Performance comparison between the case-control approach with five control per case and the divide-and-conquer approach with three subsets using the Meta method to combine. Estimated bias and empirical 95% credible interval coverage simulation study results by factors and methods. Error bars show 95% confidence intervals for the estimated bias and coverage.

We also compare computing time for the different methods, with results shown in Fig 7. For the full-data analysis of benchmark datasets with 60,000 posterior samples, the runtime was 6.25 days when the clustering proportion is 0.0009 (dark), whereas the runtime of the benchmark dataset (0.002) reached the HPC 7-day wall-time limit and was terminated at 90% progress. Based on the observed progress up to termination, the total runtime was projected to be 7.78 days (light). We therefore used 35,000 samples for this benchmark dataset to ensure completion within the wall-time limit; it took 3.99 days to be completed. This suggests that the higher clustering proportion contributes to a longer runtime. Because all subsampling datasets are likewise based on 60,000 samples, we use the estimated runtime of 7.78 days for that benchmark dataset (0.002). In general, both strategies show a marked improvement in terms of the total computation time. The results further show that datasets with a larger clustering proportion require a longer runtime when using the case-control strategy. This relationship is not observed for the divide- and-conquer strategy. In other words, computation time for the case-control approach appears to be driven by the number of 1’s pairs, while computation time for the divide-and-conquer approach mainly depends on the subsample size in each subdivision.

**Fig 7.**
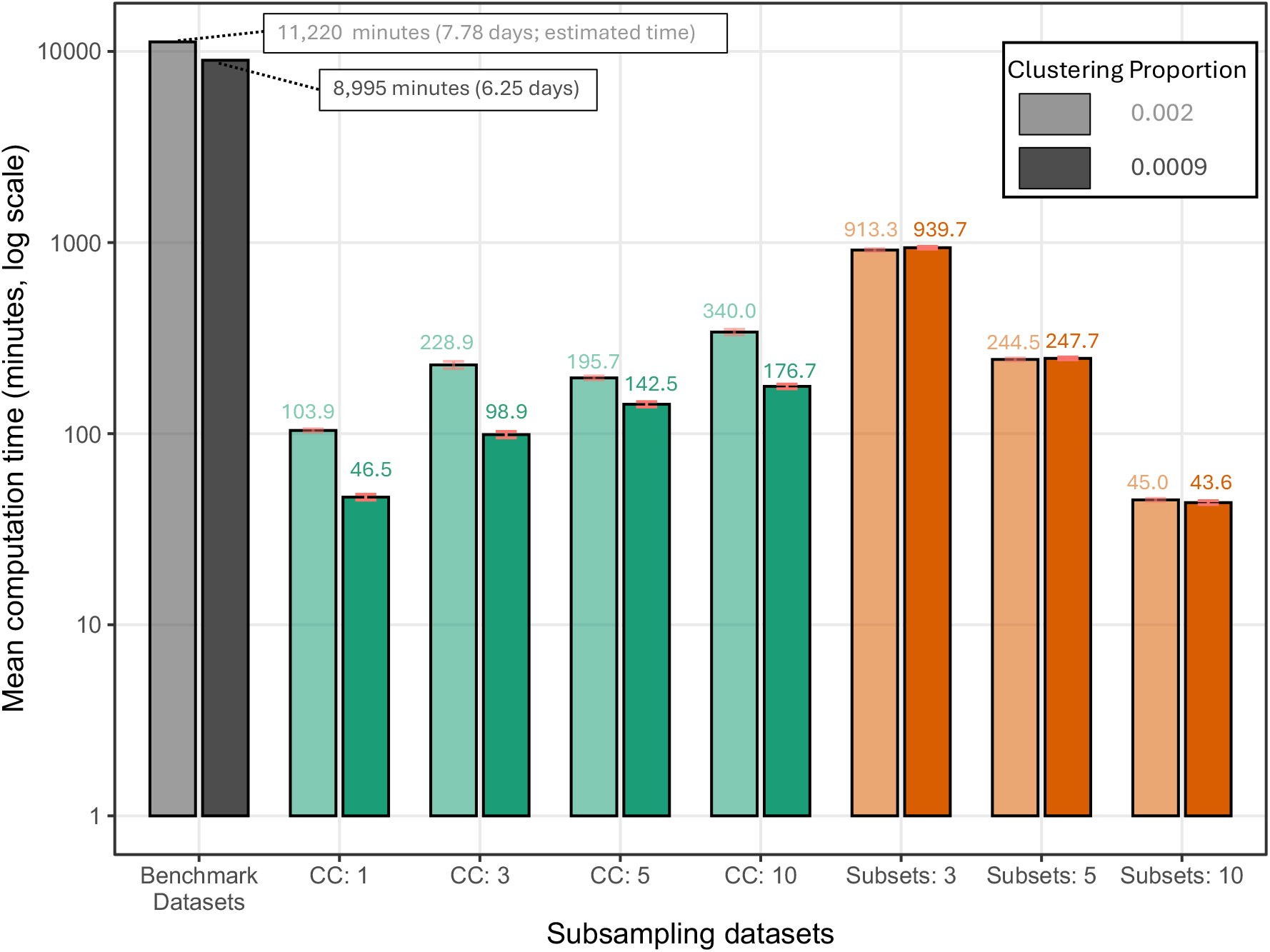
Computation time by subsampling dataset and methods, with average runtime labeled (in minutes). Light-colored bars indicate a clustering proportion of 0.002; dark-colored bars indicate a clustering proportion of 0.0009. CC denotes the case-control approach; for example, CC:1 indicates that one control was selected per case. For the divide-and-conquer approach, computation time was measured as the sum of runtime across all subdivisions and thus does not reflect potential gains from parallel execution. Note: All simulations used 60,000 MCMC samples. For the ground truth, 60,000 samples were used at clustering proportion 0.0009, while the 0.002 run did not finish within 7 days; thus, 35,000 samples were used instead, and 7 days is shown for comparison in the figure.

Based on the results from two strategies, we conclude that the case-control approach generally outperforms the divide-and-conquer approach. For both pairwise clustering proportions, using at least three controls per case seems to provide a good balance of performance and efficient use of computational resources.

## 4. Mycobacterium tuberculosis in Kaohsiung, Taiwan

We apply the case-control approach for pairwise analysis to the full dataset which includes 4,154 individuals with Mtb in Kaohsiung, Taiwan. We choose three controls per case and note that the pairwise clustering proportion of the full dataset is 0.0006, including n=5,453 1’s pairs and n=16,359 0’s pairs formed from n=4,152 individuals. We collect 5,000 samples from the joint posterior distribution after removing 10,000 iterations prior to convergence of the algorithm and thinning the remaining samples by a factor of 5 to reduce correlation. We perform the analysis on an Apple M3 system with 16 GB of memory, and the full computation requires nearly 10 days (14123.49 mins) to complete. The computation time of the full dataset without sampling is unknown, but we expect it to be substantially longer than this based on the results we show in Fig 7. We find that smaller age difference between the paired individuals (OR = 0.92 for every two-year increase in age difference, 95% CrI: 0.91-0.94), younger total age (OR = 0.86 for every two-year increase in the combined age, 95% CrI: 0.85-0.88), and closer geographic proximity (OR = 0.59 for every 5 km increase in spatial distance, 95% CrI: 0.54-0.64) were associated increased odds of being in the same genomic cluster.

To validate the results, we also apply the divide-and-conquer approach using the Meta method to combine posterior results across subsets for the whole dataset. We estimate standardized pairwise covariates from the full dataset and then randomly divided the full dataset into six subsets (around 630 cases for each subset) to conduct pairwise analysis. The choice of subset size is based on our maximum available computational capacity. For each subset, we collect 5,000 samples from the joint posterior distribution after removing 10,000 iterations prior to convergence of the algorithm and thinning the remaining samples by a factor of 5 to reduce correlation. We ran these pairwise analyses on six subsets in parallel using HPC clusters on either Ice Lake or Cascade Lake CPUs with 200 GB of memory. Each subset took around four days. Using this approach, we find that smaller age difference between the paired individuals (OR = 0.94 for every two-year increase in age difference, 95% CrI: 0.93-0.95), younger age (OR = 0.89 for every two-year increase in the combined age, 95% CrI: 0.88-0.90), and closer geographic proximity (OR = 0.70 for every 5 km increase in spatial distance, 95% CrI: 0.66-0.74) were associated increased odds of being in the same genomic cluster.

We compared results from two strategies in Fig 8. Overall, the results showed overall similar patterns. The posterior estimates of two age-related non-spatial factors were closely aligned, whereas the estimate of spatial distance showed a larger discrepancy. Such differences were consistent with our simulation results. Given the poorer bias performance observed in the simulation study, the case-control approach provides more reliable estimates.

**Fig 8.**
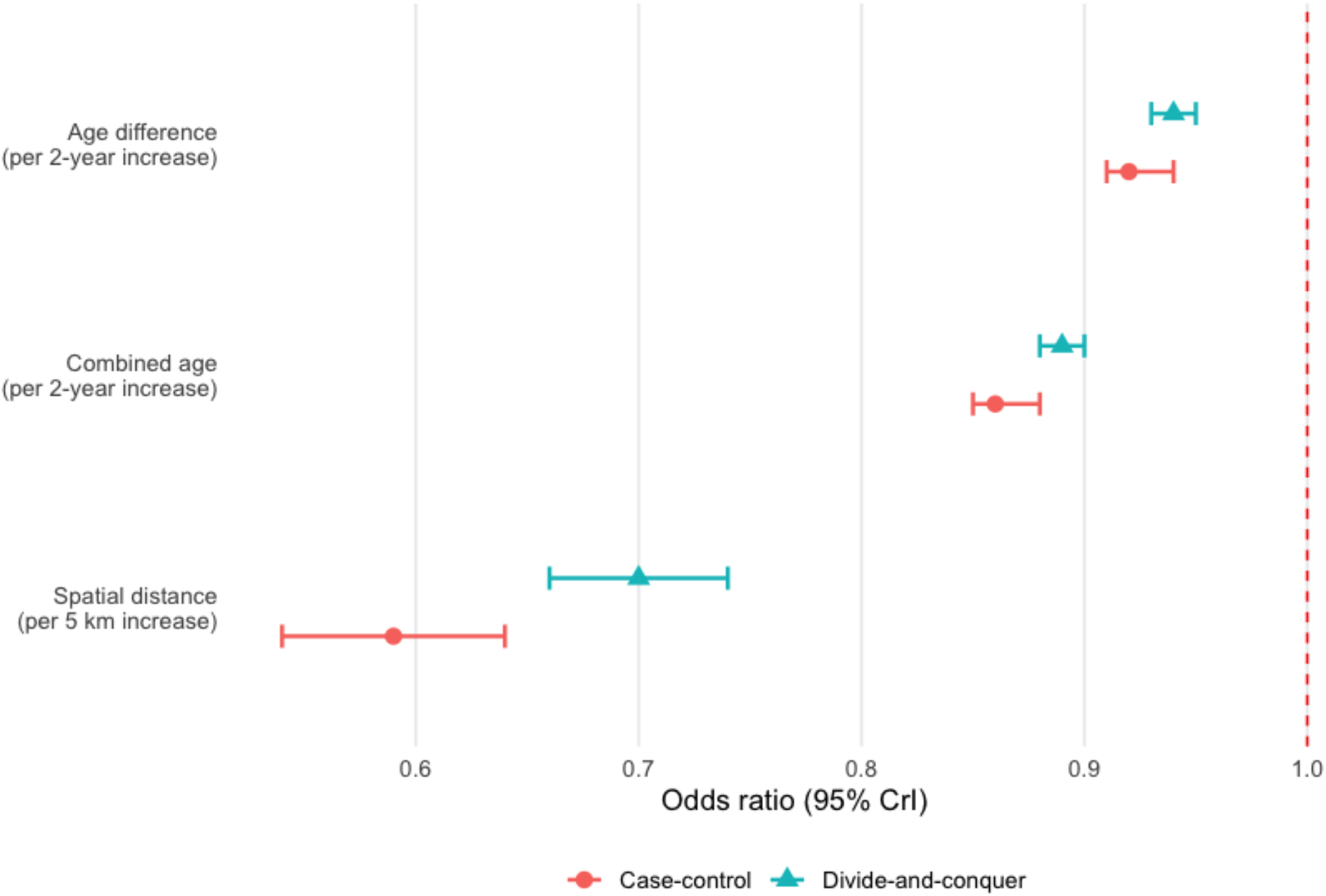
Comparison of odds ratio posterior means and 95% credible intervals for predictors on the full dataset using the case-control and divide-and-conquer methods.

## 5. Conclusion and Discussion

In this study, we compare case-control and divide-and-conquer strategies to address computational challenges for pairwise analysis of genomic and spatial data in a spatial Bayesian setting. We evaluate these methods in term of the bias, coverage and length of the credible intervals, and computation time. Based on our results in this data setting, we suggest using the case-control method with at least three controls per case and increasing the number of controls per case if the pairwise clustering proportion is much lower than 0.0009.

We also observe that the case-control approach performs better than divide-and-conquer for estimates related to the spatial covariate, while both approaches produced similar non-spatial estimates (e.g., age difference) across subdivision levels (Fig 4 – 6). As we increased the number of subdivisions, the spatial distance estimates performed worse, with wider bias distributions and lower coverage, and the spatial factor also exhibits a consistently negative bias across subdivision levels. We find that this issue is driven by the large effect size rather than the spatial nature of covariate (Fig S4). Accordingly, we suggest the case-control method is preferred as it appears more robust to the effect sizes associated with covariates of interest.

One possible explanation for the poorer performance of the divide-and-conquer approach is that partitioning the dataset across subsets effectively removes dyads spanning different subsets from the analysis. Because clustered dyads are extremely sparse, loss of these informative positive pairs may disproportionately weaken the transmission signal and contribute to attenuation of effect estimates toward the null. In contrast, the case-control approach retains all clustered dyads while subsampling only non-clustered pairs, thereby preserving the primary informative signal underlying the exposure–outcome relationship. This may explain its improved precision and reduced bias relative to the divide-and-conquer approach.

Future work should investigate alternative analytic approaches that can balance pair retention, computational efficiency, and bias reduction across a wider range of data structures and outcome types. The outcome of the model we investigated in this study is binary, while other pairwise outcomes such as SNP distance, patristic distance, or probability of transmission may also be of interest. In these cases, the issue of loss unique pairs after subsampling are not relevant, but other challenges are introduced–––such as the selection criteria for pairs with a large genetic distance (i.e., those not possibly associated with transmission) when subsampling the full dataset.

Pairwise analysis plays a critical role in understanding transmission, as it can uncover associations between person-to-person transmission. Bayesian network and spatial methods represent powerful tools in this setting but are computationally demanding. Our study identifies computationally tractable alternatives to a full dataset analysis that can preserve inference for the primary regression parameters of interest. Building on our approach to large paired genomic and spatial data, future studies should explore novel ways of pairwise analysis on TB and other infectious diseases, including pairwise pathogen, individual, and environmental risk factors.

## Supporting information captions

**S1 Fig. Mean 95% credible interval width and the corresponding 95% confidence intervals by method and factor using the case-control approach**.

**S2 Fig. Estimated bias and empirical 95% credible interval coverage simulation study results by factors and methods using the divide-and-conquer approach**.

**S3 Fig. Mean 95% credible interval width and the corresponding 95% confidence intervals for the mean by method and factors using the divide-and-conquer approach**.

**S4 Fig. Estimated bias and empirical 95% credible interval coverage simulation study results (0.0009) by factors and methods after swapping the effect sizes of age difference and spatial distance using the divide-and-conquer approach**.

**S5 Fig. Comparison of mean 95% credible interval width and the corresponding 95% confidence intervals between the case-control approach with five control per case and the divide-and-conquer approach with three subsets using the Meta method to combine**.

## Author contributions

**Conceptualization:** Yu Lan, Ted Cohen, Joshua L. Warren.

**Data curation:** Chieh-Yin Wu, Hsien-Ho Lin.

**Funding acquisition:** Hsien-Ho Lin, Ted Cohen.

**Investigation:** Yu Lan.

**Methodology:** Yu Lan, Ted Cohen, Joshua L. Warren.

**Supervision:** Ted Cohen, Joshua L. Warren.

**Visualization:** Yu Lan.

**Writing – original draft:** Yu Lan.

**Writing – review & editing:** Yu Lan, Chieh-Yin Wu, Hsien-Ho Lin, Ted Cohen, Joshua L. Warren.

## References

1. Bryant JM, Schürch AC, Van Deutekom H, Harris SR, De Beer JL, De Jager V, et al. Inferring patient to patient transmission of Mycobacterium tuberculosis from whole genome sequencing data. BMC infectious diseases. 2013;13:1–12.

2. Stimson J, Gardy J, Mathema B, Crudu V, Cohen T, Colijn C. Beyond the SNP threshold: identifying outbreak clusters using inferred transmissions. Molecular biology and evolution. 2019;36(3):587–603.

3. Didelot X, Kendall M, Xu Y, White PJ, McCarthy N. Genomic epidemiology analysis of infectious disease outbreaks using TransPhylo. Current protocols. 2021;1(2):e60.

4. Campbell F, Didelot X, Fitzjohn R, Ferguson N, Cori A, Jombart T. outbreaker2: a modular platform for outbreak reconstruction. BMC bioinformatics. 2018;19(Suppl 11):363.

5. Lan Y, Rancu I, Chitwood MH, Sobkowiak B, Nyhan K, Lin H-H, et al. Integrating genomic and spatial analyses to describe tuberculosis transmission: a scoping review. The Lancet Microbe. 2025.

6. Warren JL, Chitwood MH, Sobkowiak B, Colijn C, Cohen T. Spatial modeling of Mycobacterium tuberculosis transmission with dyadic genetic relatedness data. Biometrics. 2023;79(4):3650–63.

7. Fithian W, Hastie T. Local case-control sampling: Efficient subsampling in imbalanced data sets. Annals of statistics. 2014;42(5):1693.

8. Breslow NE. Statistics in epidemiology: the case-control study. Journal of the American Statistical Association. 1996;91(433):14–28.

9. Wang L, Williams ML, Chen Y, Chen J. Novel two-phase sampling designs for studying binary outcomes. Biometrics. 2020;76(1):210–23.

10. Wang H, Zhu R, Ma P. Optimal subsampling for large sample logistic regression. Journal of the American Statistical Association. 2018;113(522):829–44.

11. Wang J, Zou J, Wang H. Sampling with replacement vs Poisson sampling: a comparative study in optimal subsampling. IEEE Transactions on Information Theory. 2022;68(10):6605–30.

12. Wang C, Srivastava S. Divide-and-conquer Bayesian inference in hidden Markov models. Electronic Journal of Statistics. 2023;17(1):895–947.

13. Guhaniyogi R, Banerjee S. Meta-kriging: Scalable Bayesian modeling and inference for massive spatial datasets. Technometrics. 2018;60(4):430–44.

14. Orozco-Acosta E, Adin A, Ugarte MD. Scalable Bayesian modelling for smoothing disease risks in large spatial data sets using INLA. Spatial Statistics. 2021;41:100496.

15. Scott SL, Blocker AW, Bonassi FV, Chipman HA, George EI, McCulloch RE. Bayes and big data: The consensus Monte Carlo algorithm. Big Data and Information Theory: Routledge; 2022. p. 8–18.

16. Neiswanger W, Wang C, Xing E. Asymptotically exact, embarrassingly parallel MCMC. arXiv preprint arXiv:13114780. 2013.

17. Wu C-Y, Chen Y-A, Ioerger TR, Lan Y, Bai R-Y, Lee M, et al. Genomic Epidemiology of Mycobacterium tuberculosis in a Population-Based Cohort in Taiwan: Lineage-Specific Transmission and Spatial Clustering, 2019–2023 The Lancet Microbe. Forthcoming 2026.

18. Silverman BW. Density estimation for statistics and data analysis: Routledge; 2018.

19. Viechtbauer W. Conducting meta-analyses in R with the metafor package. Journal of statistical software. 2010;36:1–48.

20. Miroshnikov A, Conlon EM. ParallelMCMCcombine: an R package for Bayesian methods for big data and analytics. PloS one. 2014;9(9):e108425.

